# Subcellular compartments interplay for carbon and nitrogen allocation in *Chromera velia* and *Vitrella brassicaformis*

**DOI:** 10.1101/603555

**Authors:** Zoltán Füssy, Tereza Faitová, Miroslav Oborník

## Abstract

Endosymbioses necessitate functional cooperation of cellular compartments to avoid pathway redundancy and streamline the control of biological processes. To gain insight into the metabolic compartmentation in chromerids, phototrophic relatives to apicomplexan parasites, we prepared a reference set of proteins probably localized to mitochondria, cytosol and the plastid, taking advantage of available genomic and transcriptomic data. Training of prediction algorithms with the reference set now allows a genome-wide analysis of protein localization in *C. velia* and *V. brassicaformis*. We confirm that the chromerid plastids house enzymatic pathways needed for their maintenance and photosynthetic activity, but for carbon and nitrogen allocation, metabolite exchange is necessary with the cytosol and mitochondria. This indeed suggests that the regulatory mechanisms operate in the cytosol to control carbon metabolism based on the availability of both light and nutrients. We discuss that this arrangement is largely shared with apicomplexans and dinoflagellates, possibly stemming from a common ancestral metabolic architecture, and supports the mixotrophy of the chromerid algae.

## INTRODUCTION

Endosymbiotic organelles play crucial roles in cellular biochemistry. Mitochondria, alpha-proteobacterial endosymbionts of eukaryotes (Oborník 2019; Gruber 2019), represent an energetic hub, where balancing of catabolic and anabolic processes takes place tightly regulated with the speed of respiration (e.g. Searcy 2003; Pagliarini and Rutter 2013; Gray 2015). In plastids, domesticated cyanobacteria (Gruber 2019; Oborník 2019), inorganic carbon is fixed into sugars and several essential compounds are synthesized, such as fatty acids, isoprenoid units, tetrapyrroles, and amino acids (e.g. Tetlow et al. 2017). These semiautonomous endosymbiont-derived organelles contain genetic information and their own translation apparatuses, but by far do not encode all the proteins required for their function. Due to endosymbiotic gene transfer, most essential genes were transferred from their genomes to the nuclear genome and the organelles are greatly dependent on the import of proteins synthesized in the cytosol (e.g. Mallo et al. 2018). Sorting of proteins to subcellular locations specifically is therefore crucial for the correct function of both the proteins and the organelles, and thus targeting signals and protein translocation are key to our understanding of organellar biology (Kunze & Berger 2015).

Proteins destined to plastids and mitochondria typically encode an N-terminal motif, i.e. a targeting presequence. The targeting presequence of mitochondrial proteins is termed the mitochondrial transit peptide (mTP) and has the physicochemical properties of an amphiphilic helix. Similarly, the chloroplast transit peptides of primary algae (rhodophytes, chlorophytes and glaucophytes) and plants are amphiphilic helices, though they are typically enriched in hydroxylated amino acids and less positively charged than mTPs (Kunze & Berger 2015; Garg & Gould 2016). In comparison, complex algae (those that maintain eukaryotic endosymbionts) including the chromerids, target proteins to the plastid via the endomembrane (secretory) pathway, using chloroplast transit peptides directly preceded by an ER signal peptide, which are referred to as bipartite targeting sequences (BTS) (reviewed in Patron & Waller 2007).

Reconstruction of ancestral traits (Joy et al. 2016) allows us to unveil changes in lifestyle and genome organization in an evolutionary perspective and to compare functionalities among the organisms of interest. The discovery and genome characterization of chromerids *Chromera velia* and *Vitrella brassicaformis*, the closest known photosynthetic relatives of apicomplexan parasites, have provided an excellent framework to study the transition from free-living phototrophs to obligate parasites (Moore et al. 2008; Oborník et al. 2009; Janouškovec et al. 2010; Burki et al. 2012; Janouškovec et al. 2015; Woo et al. 2015; Füssy & Oborník 2017b). Much knowledge has accumulated about the function of the apicomplexan remnant plastid, the apicoplast (reviewed in Boucher et al. 2018), which given their shared origin structurally and molecularly resembles the photosynthetic plastid of chromerids (Moore et al. 2008; Janouškovec et al. 2010; Woo et al. 2015). Nevertheless, the protein composition of the chromerid plastid is largely unknown, except for a recent work that focused on *C. velia* photosystems (Sobotka et al. 2017), and therefore a pre-transition model of the apicoplast could not be studied in detail. The mitochondrial genome of apicomplexans is massively reduced in gene content and found to contain only three protein-coding genes, *cox1, cox3* and *cyb*, with the majority of mitochondrial proteins requiring import from cytosol (Nash et al. 2008; Janouškovec et al. 2013; Flegontov et al. 2015). Strikingly, the *C. velia* mitochondrion was found to contain only two of these genes, *cox1* and *cox3* (Flegontov et al. 2015). It has been hypothesized that the reduction of the apicomplexan (and chromerid) mitochondrial genome could be linked with the change in lifestyle strategy, particularly a change to facultative anaerobiosis (Dorrell et al. 2013), which is consistent with the observed reduction of the respiratory chain in all myzozoans (dinoflagellates, chromerids and apicomplexans) (Flegontov et al. 2015; Oborník & Lukeš 2015).

The relatively small nuclear genome size (up to 193 Mb) and largely complete sequence data of chromerids (Woo et al. 2015) make them ideal for large-scale targeting signal recognition and, by extension, organellar proteome prediction. Up to now, plastid proteomes have been determined in only a handful of organisms, mainly plants and green algae (van Wijk & Baginsky 2011; Terashima et al. 2011; Dorrell et al. 2017), but also a handful of complex algae and protist parasites (Hopkins et al. 2012; Boucher et al. 2018). Similarly, mitochondrial proteomic data are rather scarce and focused on model organisms, such as humans (Calvo & Mootha 2010; Palmfeldt & Bross 2017), yeast (Gonczarowska-Jorge et al. 2017), plants (Huang et al. 2013), and protists (Smith et al. 2007; Atteia et al. 2009; Panigrahi et al. 2009; Danne et al. 2013; Gawryluk et al. 2014).

The aim of the work is to define and characterize the subcellular proteomes of chromerids by bioinformatic tools with an emphasis on plastid- and mitochondrion-destined proteins. For the analysis, we compiled sets of compartment-specific proteins of *C. velia* and *V. brassicaformis* and optimized the performance of the ASAFind (Gruber et al. 2015) prediction tool with the reference amino acid frequency matrices. Our analyses bring first implications on carbon and nitrogen allocation among the plastid, cytosol and mitochondria in chromerids, suggesting interplay of these compartments is in place for efficient carbon metabolism under changing light and nutrient conditions. This work also confirms biochemical peculiarities ancestrally shared with apicomplexans and dinoflagellates, such as the lack of the canonical mitochondrial pyruvate decarboxylase and cytosolic amino acid synthesis.

## DATA SOURCES AND METHODS

The sequence data of the chromerid algae *Chromera velia* CCMP2878 and *Vitrella brassicaformis* CCMP3155 were retrieved from CryptoDB (www.cryptodb.org, version 34). Additional transcriptomic data were retrieved from NCBI GenBank (Dorrell et al. 2014; Woehle et al. 2011) and MMETSP sequence databases (MMETSP0290 and MMETSP1451, (Keeling et al. 2014; Cohen et al. 2016). The sequence data were annotated using the information available at KEGG servers (Kanehisa et al. 2017) and using the InterProScan annotation tool of Geneious (Jones et al. 2014; Kearse et al. 2012).

Plastid-targeted reference sequences were identified based on several lines of evidence; a) the protein had a clear role in the plastid metabolism (in synthesis of pigments and cofactors, or being a subunit of the photosynthetic machinery, etc.), with an emphasis on filling the gaps between well-defined enzymatic steps; b) reassuring sequence completeness, an N-terminal extension (40-80 amino acids) that could possibly encode a bipartite targeting sequence preceded the mature protein (as determined by InterProScan), though essentially no targeting sequence prediction was employed to avoid circular reasoning (including predictor-positive proteins and using them to evaluate this predictor); c) there was a phylogenetic relationship to another plastid-targeted protein among chromerids or Apicomplexa (in case of ribosomal proteins; Gupta et al. 2014). Mitochondrial references were compiled similarly, only the N-terminal extension was found shorter. Cytosolic references lacked an extension and secretory proteins had an identifiable role in the endomembrane system or at the cytoplasmic membrane. Metabolic gaps were filled by targeted BLAST searches in the genomic (CryptoDB) as well as transcriptomic data (CryptoDB, GenBank, MMETSP) using known-function apicomplexan sequences and KEGG orthologs as queries.

To define the best tool for the subcellular localization of proteins, the sets of reference sequences of *Chromera* and *Vitrella* were analyzed by prediction algorithms. The tools tested were selected to be suitable for large-scale analyses and included: TargetP (Emanuelsson et al. 2000), SignalP (v. 4.1) (Petersen et al. 2011), ASAFind (Gruber et al. 2015), HECTAR v1.3 (Gschloessl et al. 2008), MultiLoc2 (Blum et al. 2009), PrediSi (Hiller et al. 2004) and PredSL (Petsalaki et al. 2006). All the prediction algorithms except HECTAR were run locally with default parameters; SignalP was run with sensitive cutoff values (-u 0.3 -U 0.3). The sensitivity (proportion of recognized true positives) and precision (proportion of positive results, also termed the positive predictive value) of the prediction algorithms were compared, setting a certain threshold specified for each of the predictors. Sensitivity was computed as:

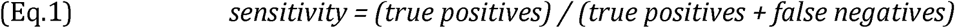

value was computed as:

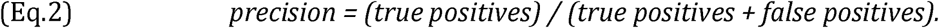

An optimal threshold would cover maximum positive proteins while including a low number of false positives (proteins falsely predicted to the organelle in question).

Bit score-weighted matrices of amino acid positions surrounding the signal cleavage sites were calculated separately for *C. velia* and *V. brassicaformis*, as described by Crooks et al. (2004). Only cleavage sites which were consensually (by majority agreement) predicted by PrediSi, PredSL, SignalP and ASAFind were considered. The transit peptide sequence logos and frequency plots of plastid-targeted proteins from *C. velia* and *V. brassicaformis* were created with WebLogo (Crooks et al. 2004; http://weblogo.berkeley.edu/; version 2.8.2).

Where applicable, closest hits for proteins were found using BLAST against nr or RefSeq databases, and using DIAMOND (Buchfink et al. 2014) against an in-house made database consisting from sequences collected from NCBI, MMETSP (Keeling et al. 2014; Cohen et al. 2016) and Ensembl Genomes (release 37; Kersey et al. 2016). Sequences were aligned using MAFFT v.7 (Katoh & Standley 2013) and automatically trimmed by trimAL (Capella-Gutiérrez et al. 2009). Maximum likelihood trees were inferred from the trimmed alignments using the best-fitting substitution model as determined by the IQ-TREE –m TEST option limited to LG matrix by -mset (Nguyen et al. 2015). Branch supports were determined by rapid bootstrapping followed by 1,000 ultra-fast bootstrap replicates (-bb 1000).

## RESULTS

### Sequence completeness and reference compilation

During the annotation and alignment of reference proteins, we noticed that some contigs retrieved from CryptoDB (Woo et al. 2015) are apparently truncated at their 5’-ends. The sequence data appear gene-rich but are still highly fragmented, with 5,966 and 1,064 genomic scaffolds assembled for *C. velia* and *V. brassicaformis*, respectively, although the number of chromosomes in *C. velia* is estimated to be much smaller (Vazac et al. 2017). Since it is essential to identify a protein’s complete N-terminus to predict its localization to the plastid or to mitochondria (see Introduction), we pursued an independent assessment of N-termini completeness. We searched for (almost) identical transcripts in chromerid transcriptomes available in GenBank and MMETSP and used these contigs to extend those from CryptoDB towards their 5’-end, where possible. While these fused contigs are indeed chimeras of orthologs from different strains (the transcripts were not completely identical), we assume they code for *bona fide* N-termini in all these sequenced strains of *C. velia* and *V. brassicaformis*. These chimeric contigs are for clear distinction marked in the reference sequence list (Supplementary Table S1). Other, apparently truncated contigs were omitted from the reference sets.

We performed a systematic search of housekeeping and metabolically active proteins to obtain the reference sets from *C. velia* and *V. brassicaformis* data. To find plastid references, we searched for pathways intimately linked with their biogenesis and photosynthesis, including the biosynthesis of fatty acids (type II FAS) and lipids, iron-sulfur clusters, terpenoids and terpenoid derivatives (photosynthetic pigments, vitamins), tetrapyrrole cofactors, protein translocator subunits and components of the photosynthetic electron transport chain (recently characterized by Sobotka et al. 2017) (Fig. 1), and the enzymes of carbon fixation (Fig. 2). Among mitochondrial pathways, we looked for the enzymes of the tricarboxylic acid (TCA) cycle and the components of the respiratory chain (Flegontov et al. 2015) (Fig. 2). From pathways localized in the cytosol, we identified the enzymes of glycolysis, storage amylopectin biosynthesis and breakdown (Coppin et al. 2005), and the enzymes of the pentose phosphate cycle that we distinguished from the enzymes of the plastid carbon fixation by their N-terminal extensions (Fig. 2). Using phylogenetic analyses (not shown), we also identified ribosomal proteins from all three translationally active compartments (Gupta et al. 2014). We also included biosynthesis pathways of several amino acids and other compounds, where enzymatic steps showed consistent localization (Fig. 2). Organellar genome-encoded proteins were dropped from the reference sets (Supplementary Table 1). In a scarcity of experimental data, we believe this is the most reliable approach to compile reference sequences.

**Figure 1:**
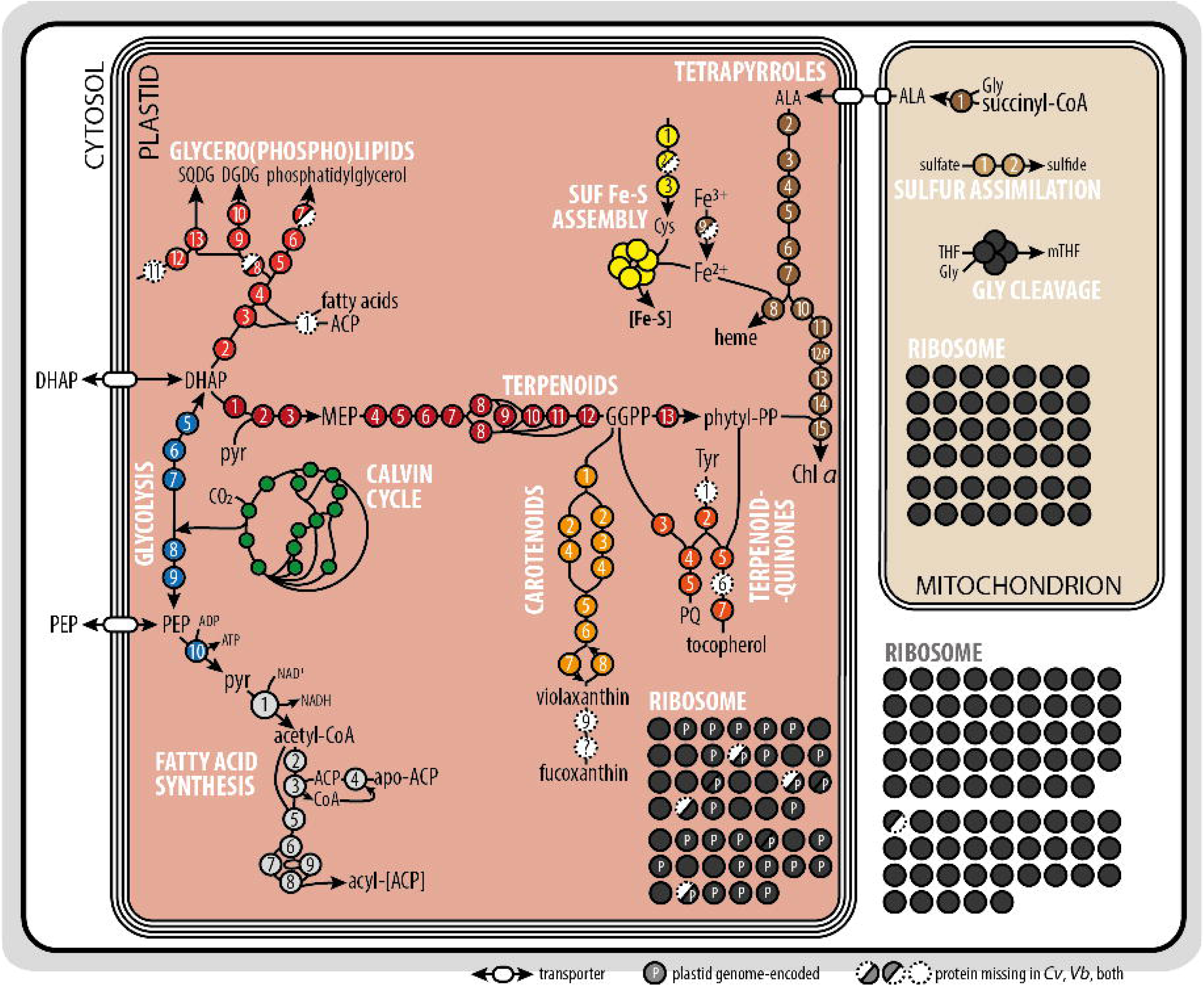
**Overview of reference pathways**, focused on the inter-connected reactions of terpenoid, lipid and tetrapyrrole biosynthesis, mitochondrial sulfur assimilation and glycine cleavage, and ribosomal proteins. Pathways are color-coded and enzymes/proteins are numbered according to the Supplementary Table S1. Abbreviations: ACP – acyl-carrier protein, ALA – delta-aminolevulinic acid, Chl *a* – chlorophyll *a*, DGDG – digalactosyldiacylglycerolipids, DHAP – dihydroxyacetone phosphate, GGPP – geranylgeranyl pyrophosphate, IMS – inter-membrane space, MEP – methyl-D-erythritol 4-phosphate, PEP – phospho*enol*pyruvate, pyr – pyruvate, SQDG – sulfoquinovosyldiacylglycerolipids, THF – tetrahydrofolate.

**Figure 2:**
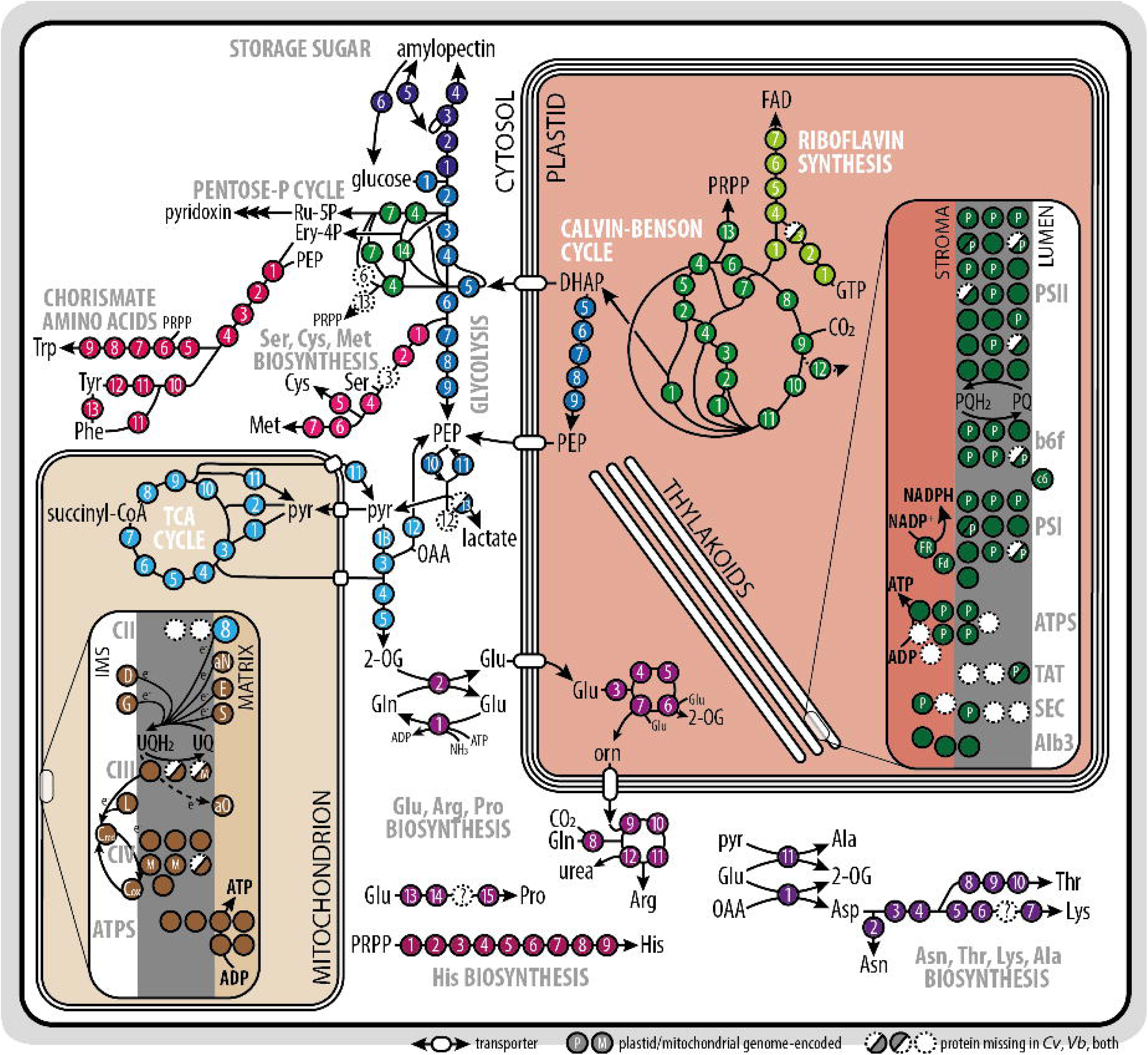
**Overview of reference pathways, continued**, focused on the inter-connected reactions of carbohydrate and amino acid biosynthesis, mitochondrial respiration and plastid photosynthesis. Pathways are color-coded and enzymes/proteins are numbered or coded according to the Supplementary Table S1. 2-OG – 2-oxoglutarate, DHAP – dihydroxyacetone phosphate, Ery-4P – erythrose 4-phosphate, FAD – flavin adenine dinucleotide, IMS – inter-membrane space, OAA – oxaloacetate, orn – ornithine, PEP – phospho*enol*pyruvate, PQ – plastoquinol, PRPP – 5-phosphoribosyl-1-pyrophosphate, pyr – pyruvate, Ru-5P – ribulose 5-phosphate, TCA – tricarboxylic acid, THF – tetrahydrofolate, UQ – ubiquinone.

Though most of the searched pathways are near-complete, we failed to identify representatives of some enzymatic steps in the available transcriptomic and genomic data. These include the acyl-[acyl carrier protein (ACP)] ligase (in both chromerids) and two lipid phosphatases of the glycero(phospho)lipid biosynthesis (phosphatidate phosphatase in *C. velia*, phosphatidylglycerophosphatase in *V. brassicaformis*), several enzymes of the carotenoid (neoxanthin synthase), terpenoid-quinone (tocopherol cyclase), and amino acid biosynthesis pathways (undescribed steps of Lys and Pro synthesis), and one enzyme of the pentose phosphate cycle (cytosolic ribose 5-phosphate isomerase; Fig. 1, Fig. 2). Some of these steps are possibly catalyzed by distantly related enzymes that were not recognized by our searches, but other absences may be of biological relevance. Ribose 5-phosphate isomerase is necessary to recycle ribulose 5-phosphate, but a cytosolic isoform was not recovered, and maybe alternatively spliced transcripts encode for a truncated, non-plastid-targeted protein. Similarly, the final steps of fucoxanthin biosynthesis are missing. Although there are biochemical data that *Chromera* and *Vitrella*, respectively, accumulate (iso)fucoxanthin and vaucheriaxanthin – both derivates of neoxanthin – as accessory photosynthetic pigments (Moore et al. 2008; Oborník et al. 2012), neoxanthin synthase was not found in our data, while the last enzymes of the respective pathways are unknown (Mikami & Hosokawa 2013). Plant neoxanthin synthase is a neofunctionalized lycopene cyclase (Bouvier et al. 2000), opening the possibility that a promiscuous activity of the latter enzyme is responsible for neoxanthin synthesis in chromerids. In comparison, the absence of acyl-ACP synthase might be compensated if fatty acids are not released from the bond with the ACP but rather directly used for lipid synthesis (Bisanz et al. 2006). Alternatively, intermediate glycerolipids (diacylglycerol esterified with fatty acids of various length and saturation) could be imported from other cell compartments (Jouhet et al. 2007).

### Prediction performance

ASAFind and HECTAR (Gruber et al. 2015; Gschloessl et al. 2008) were designed to predict the BTS of plastid proteins in complex algae, particularly in heterokonts. ASAFind identifies plastid proteins based on the output of SignalP (Petersen et al. 2011) and a sliding-window scan for the highly conserved Phe residue around the predicted cleavage site. HECTAR uses a combination of predictors in three decision modules and aims to classify proteins based on presence of four types of targeting modules: signal peptides, type II signal anchors, chloroplast transit peptides, and mitochondrial transit peptides. To predict the mTP, several predictors are available. These include TargetP, MultiLoc2, and HECTAR (Emanuelsson et al. 2000; Blum et al. 2009; Gschloessl et al. 2008).

Both ASAFind and HECTAR were specifically trained on stramenopile sequences, though ASAFind offers the possibility to use an alternative bit score matrix derived from a training set. TargetP and MultiLoc2 were trained using plant and animal sequences. Having reference dataset at hand, we could test the performance of these algorithms on chromerid datasets, which has been unknown. Our results are summarized in Table 1 (full results in Supplementary Table S2). Mitochondrial predictors offered only moderate sensitivity and precision, with *V. brassicaformis* sequences being better resolved. We defined two thresholds for each predictor, one having higher sensitivity (around 75 %), the other more selective (with around 85 % precision). HECTAR offers higher precision with the 75%-level sensitivity compared to other mitochondrial predictors (Supplementary Fig. S3). With plastid sequences, ASAFind and HECTAR performed comparably, having both high sensitivity and precision. However, it must be noted that the threshold for HECTAR is very low and most of our plastid controls were marked as “secretory proteins” by this predictor. Our datasets might be over-fit by lacking enough endomembrane system proteins as negative controls, therefore precision is expected to drop with broader reference sets. We also used our plastid references to derive bit score-weighted matrices of amino acids flanking the cleavage site independently for *C. velia* and *V. brassicaformis* (as in Gruber et al. 2015). Using this matrix, we could further improve the performance of ASAFind (Table 1). This suggests that indeed chromerid plastid-targeting presequences differ from those of stramenopiles, but possibly also from each other.

**Table 1:** Performance of various localization predictors as employed in this study. ASAFind and HECTAR-plastid were used for plastid reference assessment. Note that ASAFind has a cumulative score, with our data reaching up to values of 6. ASAFind+ denotes ASAFind with species-specific bit score matrices designed with *C. velia* and *V. brassicaformis* plastid references. TargetP, HECTAR-mitochondrion and MultiLoc2 were used for mitochondrial predictions and two thresholds are presented for these predictors, one aimed at higher sensitivity (s – 75%), the other aimed at high precision (p – 85%).

|  | ASAFind | ASAFind+ | HECTAR | TargetP - mTP |  | HECTAR - mTP |  | MultiLoc2 - mTP |  |
| --- | --- | --- | --- | --- | --- | --- | --- | --- | --- |
|  |  |  |  | (s) | (p) | (s) | (p) | (s) | (p) |
| *C. velia* |  |  |  |  |  |  |  |  |  |
| THRESHOLD: | 1.4 | 1.4 | 0.05 | 0.35 | 0.8 | 0.14 | 0.41 | 0.25 | 0.95 |
| SENSITIVITY: | 0.774 | 0.830 | 0.769 | 0.757 | 0.586 | 0.748 | 0.586 | 0.748 | 0.361 |
| PRECISION: | 0.921 | 0.921 | 0.872 | 0.375 | 0.844 | 0.529 | 0.855 | 0.485 | 0.736 |
| *V. brassicaformis* |  |  |  |  |  |  |  |  |  |
| THRESHOLD: | 1 | 1 | 0.16 | 0.56 | 0.81 | 0.18 | 0.37 | 0.64 | 0.89 |
| SENSITIVITY: | 0.759 | 0.897 | 0.744 | 0.757 | 0.586 | 0.748 | 0.559 | 0.755 | 0.582 |
| PRECISION: | 0.886 | 0.916 | 0.967 | 0.712 | 0.855 | 0.664 | 0.861 | 0.697 | 0.842 |

To visualize the amino acid enrichment around the signal cleavage site, we created logo-plots of amino acid frequencies in this motif in chromerids. We found that both chromerids have a conserved Phe residue just after the cleavage site, though in *C. velia* the Phe motif is more frequently found (Fig. 3).

**Figure 3:**
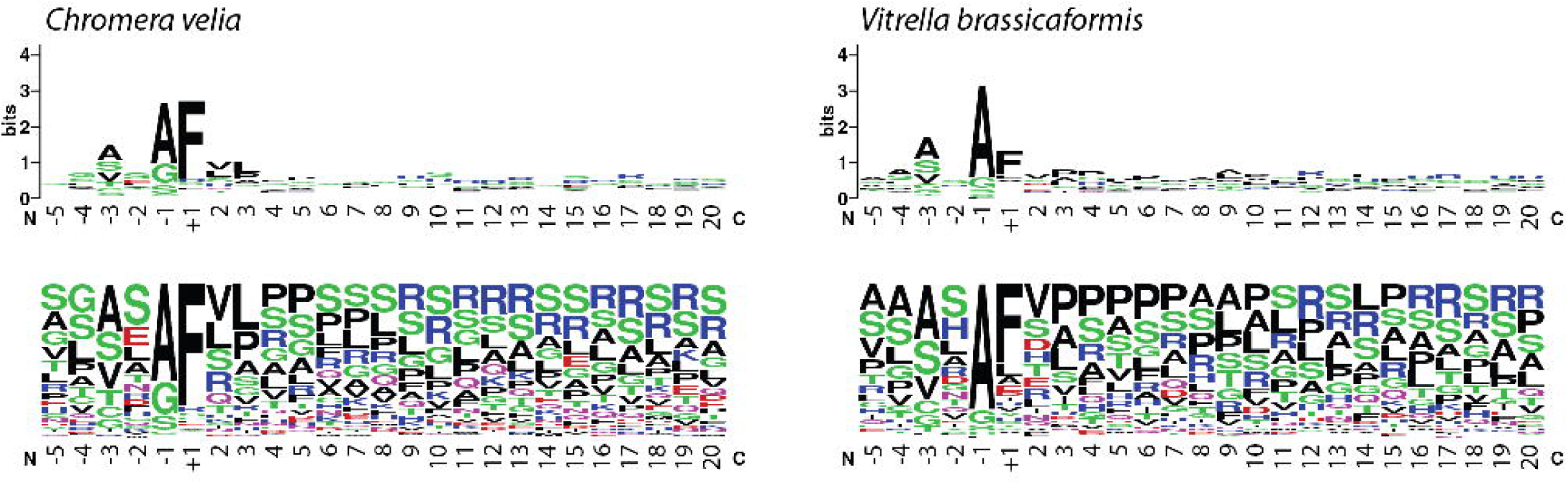
Sequence motifs surrounding the signal cleavage site differ in chromerids. While *C. velia* retains a highly conserved Phe residue that follows after the cleavage site (marked by +1), in *V. brassicaformis* Phe seems less conserved. Logoplot created by WebLogo (Crooks et al. 2004).

### Plastid housekeeping

Most of the enzymes we expected to localize in chromerid plastids are directly or indirectly related to photosynthesis. These include the photosystems core subunits, the proteins of light-harvesting antennae and the photosynthetic electron transport chain, but also insertase proteins TAT, SEC and SRP/Alb3 that embed these factors in the thylakoid membrane (Fig. 2, discussed in further detail by Sobotka et al. 2017). Furthermore, we detected the many enzymatic steps that produce photosystems cofactors and accessory pigments (Fig. 1). Tetrapyrrole synthesis is one of the prime biochemical pathways of plastids, since heme is a vital component to the electron transport chain and retrograde plastid signaling, while chlorophyll is a fundamental cofactor in light harvesting complexes (de Souza et al., 2017). In *C. velia*, the tetrapyrrole pathway starts curiously with the delta-aminolevulinic acid (ALA) synthesis from glycine and succinyl-CoA in the mitochondrion, but the rest of the pathway is predicted to reside in the plastid (Kořený et al. 2011). This feature is shared by *V. brassicaformis* and indeed supported by our results (Fig. 1). Heme is also required for the function of cytosolic and mitochondrial hemoproteins (e.g., respiratory chain components), so it is exported from the chromerid plastid, as there is apparently no tetrapyrrole synthesis activity in other compartments besides ALA synthesis.

Chlorophyll *a* (Chl *a*) is the only chlorophyll species employed by chromerids (Moore et al. 2008). During the last step of Chl *a* synthesis, a terpenoid (phytyl) moiety is attached to the Mg^2+^-coordinated tetrapyrrole, chlorophyllide *a*. Phytyl diphosphate is synthesized from methyl-D-erythritol 4-phosphate (MEP) and the pathway also resides in the chromerid plastid. The alternative cytosolic (mevalonate) pathway for terpenoid biosynthesis is missing, but terpenoid diphosphates are apparently exported from the plastid for further elongation by cytosolic and mitochondrial polyprenyl diphosphate synthases to produce, among others, ubiquinone for the mitochondrial respiratory chain (Supplementary Table S1, Imlay & Odom 2014).

Terpenoid derivatives include carotenoids and terpenoid quinones. Carotenoids violaxanthin and (iso)fucoxanthin are accessory photosynthetic pigments involved in light-harvesting, and the de-epoxidation of violaxanthin was suggested to be fundamental for photoprotection in *C. velia* (Kotabová et al. 2011). Plastoquinol is an electron transport chain electron mediator, while tocopherol has a protective role in oxidative stress (Müller & Kappes 2007). The plastid localization of these pathways therefore conditions the photosynthetic abilities of chromerids.

Fatty acid and lipid synthesis are important for the generation of plastid membranes and the modulation of their physicochemical properties. Chromerid plastids house the type II fatty acid synthesis pathway that is fed with acetyl-CoA by the plastid pyruvate dehydrogenase complex (Foth et al. 2005). The produced acyl-ACP molecules may then be directly esterified with glycerol 3-phosphate to become intermediates of glycerolipid synthesis. Galactolipids of the mono- and digalactosyl-diacylglycerolipid groups (MGDG and DGDG) are major lipids of plastids in *C. velia* (Botté et al. 2011), probably required for proper embedding of photosystems into the thylakoid membranes but possibly also involved in plastid protein translocation (Schleiff et al. 2003). Their biosynthesis is limited to plastids in both *C. velia* and *V. brassicaformis*, as the respective protein contain plastid-targeting presequences (Fig. 1).

Flavin cofactors are critical for multiple enzymes of the TCA cycle, fatty acid oxidation, photosynthesis, respiratory chain, and *de novo* pyrimidine biosynthesis. In chromerids, riboflavin is synthesized in the plastid from a Calvin cycle intermediate, ribulose 5-phosphate, and flavin nucleotides are then supposedly exported from the plastid (Fig. 2).

Iron-sulfur clusters are components of redox proteins and in plastids, they function as the prosthetic group of the cytochrome b_6_f and the ferredoxin redox relay that signals the plastid redox state to downstream enzymes. Plastid iron-sulfur clusters are typically synthesized by the SUF system, using ferrous ions and cysteine-derived sulfur as substrates. Both Cys synthesis and SUF system are found in chromerid plastids (Fig. 1), suggesting that plastid client proteins have regular access to Fe-S clusters.

Unexpectedly, only few enzymes of amino acid biosynthesis localize to plastids in *C. velia* and *V. brassicaformis*. We could reliably predict that only three enzymes of Cys synthesis reside in the plastid, but this amino acid is directly metabolized by the SUF system to assemble Fe-S clusters. Strikingly, plant chloroplasts synthesize several amino acids (Lys, Arg, Ala, Trp, Tyr, Phe; Van Dingenen et al. 2016) and participate on nitrogen assimilation with the glutamine synthase / glutamine oxoglutarate aminotransferase (GS/GOGAT) cycle. Apparently, chromerid plastids are much less involved in nitrogen metabolism than previously studied systems (see below).

In summary, chromerid plastids are well-equipped for the biogenesis and maintenance of the photosynthetic machinery. In addition, these organelles prime fatty acid and terpenoid biosynthesis by the formation of short-chain intermediates (short-chain saturated fatty acids and monoterpenoid diphosphates), that are however exported for further processing. Heme is another crucial compound produced in this compartment. All these processes seem closely coupled to photosynthesis and, importantly, to carbon fixation by the Calvin cycle, which will be overviewed in the next section.

### Compartment interplay

Calvin cycle is the very center of the second phase of photosynthesis; it utilizes ATP and NADPH produced by the light-dependent reactions to fix carbon dioxide into carbohydrates. We identified in chromerids all the enzymes of the cycle, discriminating them from the cytosolic carbohydrate metabolism variants by their N-terminal extensions having BTS characteristics. For three molecules of CO_2_, the Calvin cycle produces one triose phosphate molecule (dihydroxyacetone phosphate – DHAP, or 3-phosphoglycerate) that can be metabolized by other pathways (see above), exported to the cytosol or reintroduced to the cycle to produce sugar phosphates with more carbons.

Triose phosphates enter the cytosol via the triose phosphate transporters, and those identified in *Plasmodium falciparum* prefer DHAP and phospho*enol*pyruvate (PEP) as substrates, while also accepting 3-phosphoglycerate (Lim et al. 2010). Upon the entry of triose phosphates to the cytosol, they can take two major paths, depending on the metabolic state. Gluconeogenetic pathway leads to the accumulation of storage saccharides (amylopectin), while glycolysis yields pyruvate, a hub compound for both anabolic and catabolic reactions. Generally, favorable and illuminated growth of algae promote storage and anabolic pathways, while dark growth and starvation promote spending of sugar phosphates in the TCA cycle and respiration.

We found enzymes involved in all processes of polysaccharide accumulation and degradation (Coppin et al. 2005). Some of the storage sugar enzymes encode signal peptides or transmembrane domains, suggestive of their extracellular or membrane-associated activity. This is consistent with chromerids forming a thick cellulose cell-wall (Moore et al. 2008; Oborník et al. 2012; Füssy & Oborník 2017a).

The Calvin cycle is not only a source of triose phosphates for cytosolic glycolysis/gluconeogenesis, but also of pentose phosphates that are the starting substrates for nucleotide (ribose 5-phosphate), amino acid (ribose 5-phosphate and erythrose 4-phosphate), and vitamin synthesis (ribulose 5-phosphate). In this respect it complements the cytosolic pentose phosphate cycle that also provides these sugar phosphates. Based on our predictions, the plastid ribulose 5-phosphate pool serves as the substrate for flavin nucleotide synthesis, while cytosolic ribulose 5-phosphate is used for pyridoxin synthesis. Notably, we could not find any cytosolic 5-phosphoribosyl-1-pyrophosphate (PRPP) synthases that would provide PRPP for the synthesis of His, Trp and nucleotides in this compartment. Two pairs of orthologs are found in chromerids. Each of these sequences has a presequence, though only two orthologs have BTS (the other pair have mTP-like N-termini and were discarded from predictions due to uncertain localization; Supplementary Table S1). Comparison with PRPP synthase sequences from apicomplexans revealed that some of those too have predicted presequences (including both *Toxoplasma gondii* ME49 paralogs), but proteins having these presequences do not cluster together (Supplementary Table S1). The localization of PRPP synthases is therefore questionable and might be achieved by alternative splicing.

Other anabolic reactions also stem from the cytosolic pool of sugar phosphates. Biosynthesis of most amino acids in chromerids depends on glycolysis and pentose phosphate cycle intermediates. Cys, Met and Ser derive from 3-phosphoglycerate, while Phe, Trp and Tyr are synthesized from erythrose 4-phosphate via chorismate, and His is a derivative of PRPP (but see above). Ala, Arg, Asp, Asn, Lys, Pro and Thr derive from 2-oxoglutarate (2-OG) transaminated to form Glu by the GS/GOGAT cycle (Fig. 2). Notably, the cytosolic 2-OG pool for GS/GOGAT cycle exists by the action of cytosolic copies of TCA cycle enzymes (Fig. 2, Supplementary Table S1). Chromerid plastids do not principally participate in the amino acid biosynthesis, with a few exceptions. A part of the Arg pathway (ornithine synthesis) is predicted to localize in the plastid, which might be a rate-limiting, regulatory measure on ornithine production via dependence on stromal ATP levels. This is not unexpected, as in plants Arg synthesis is regulated at the level of N-acetylglutamine phosphorylation (Ferrario-Méry et al. 2006). Notably, chromerids utilize argJ (Supplementary Table S1), which is a glutamate transacetylating enzyme that allows to recycle N-acetylglutamate after the production of ornithine. This reaction does not produce free acetate and thus does not require additional ATP for acetyl-CoA recycling. In *Plasmodium, Cryptosporidium* and *Eimeria*, Arg (via ornithine) is used for polyamine synthesis (Cook et al. 2007) but this pathway shows cytosolic localization congruently in apicomplexans and chromerids (Supplementary Table S1; Shanmugasundram et al. 2013). Gly may be produced by glycine hydroxymethyltransferase and threonine aldolase in mitochondria, cytosol, and plastids (Supplementary Table S1), reflecting its involvement in multiple pathways as a reaction intermediate (for plastid formylmethionine and mitochondrial ALA synthesis, for instance). There is some incongruence about the localization of Ile, Leu and Val synthesis among chromerids, as *V. brassicaformis* appears to localize Val and Ile biosynthesis to both mitochondria and plastid while Leu biosynthesis is cytosolic. In comparison, all these pathway steps are cytosolic in *C. velia*. Chromerid plastids thus appear to have minor roles in amino acid biosynthesis.

Both catabolic and anabolic reactions were found to take place in the chromerid mitochondria, though they are missing some common components. Both apicomplexans and chromerids ancestrally lack the respiratory complex I (and *C. velia* also lacks complex III) and utilize alternative NADH dehydrogenases that pass electrons to ubiquinone but do not contribute to the proton gradient (Fig. 2, Flegontov et al. 2015). Like Apicomplexa, chromerids also lack a canonical mitochondrial pyruvate dehydrogenase and instead take advantage of the promiscuous activity of the branched-chain amino acid dehydrogenase (BKCDH) (Foth et al. 2005; Danne et al. 2013; Oppenheim et al. 2014). The NAD^+^-dependent isocitrate dehydrogenase is missing and replaced by an NADP^+^-dependent isozyme, which could be linked to the loss of the canonical NADH dehydrogenase (respiratory complex I). NADP^+^-dependent isocitrate dehydrogenase might in turn support the activity of NADPH-dependent enzymes (Danne et al. 2013). This is notable because in mammals both NAD^+^ and NADP^+^ isocitrate dehydrogenases are operational, the latter typically in reverse (reductive) direction (Sazanov & Jackson 1994; Yoo et al. 2008). In contrast with apicomplexans, both fumarate hydratase types are present in chromerids (apicomplexans express only type I; Bulusu et al. 2011), while an ortholog of the conserved apicomplexan malate:quinone oxidoreductase is missing (Danne et al. 2013). Chromerids localize to mitochondria several enzymes of amino acid decomposition (Supplementary Table S1), suggesting that at least some catabolic pathways feed into the mitochondrial metabolism. A set of lactate dehydrogenases in chromerids allow fermentation of pyruvate to lactate under temporary dark anaerobic conditions (Flegontov et al. 2015; Oborník & Lukeš 2015) and participate in methylglyoxal detoxication (Cordeiro et al. 2012). Furthermore, the mitochondrial metabolism is equipped with malic enzyme (decarboxylating malate dehydrogenase), which allows the regeneration of pyruvate for anabolic reactions when the cycle is fed by fatty acid beta-oxidation. Therefore, mitochondria are well-integrated in the chromerid carbon metabolism in both catabolic and anabolic directions.

Carbon metabolism also affects the synthesis rates of nitrogen pathways. The GS/GOGAT cycle is responsible for nitrogen (ammonia) assimilation into amino acids in most phototrophs. In chromerids, GOGAT is an NADH-dependent cytosolic enzyme, and the restriction of amino acid pathways to the cytosol suggests that they are decoupled from the redox state of the plastid and rather reflect the redox state of the cytosol (Fig. 2). This also suggests that the plastids of chromerids are not as deeply involved in primary metabolism as the plastids of primary algae and plants by lacking the ability to synthesize amino acids and polysaccharides. Instead, plastid activity is crucial in lower carbohydrate and fatty acid metabolism and appears to be sensed indirectly, through the supply of triose phosphates. Under favorable conditions, the cytosol is fed with photosynthetic sugar phosphates, and reducing agents and ATP are produced by glycolysis. This promotes anabolic reactions that allow the accumulation of polysaccharides and production of amino acids and lipids. In the dark or under nutrient scarcity, the shortage of energy must be compensated by the reactions of the TCA cycle and respiration in mitochondria. Unfortunately, there are no published large-scale quantitative data suitable for tracing in more detail the metabolic flows through the described pathways.

## DISCUSSION

The physicochemical character of mitochondrial and plastid targeting presequences remains quite similar across long evolutionary distances among eukaryotes. Still, the predicting power of localization algorithms decreases with divergence between the reference and the analyzed sets. Prediction algorithms perform best if trained using lineage-specific datasets, usually based on available experimental data (e.g. (Emanuelsson et al. 2007; Kaundal et al. 2013; Gruber et al. 2015). Consistently, plastid-localization signals show specific variability among algal clades (Patron & Waller 2007). Although many tools have been implemented to determine mitochondrion- and plastid-localized proteins in *C. velia* and *V. brassicaformis* (Kořený et al. 2011; Woehle et al. 2011; Petersen et al. 2014; Flegontov et al. 2015; Woo et al. 2015; Sobotka et al. 2017), none of them have been tested on reference proteins in terms of predictive power. To find a suitable tool to predict protein localization in chromerids, we prepared a manually curated inventory of references that included proteins from plastid, mitochondrion, and several other compartments, as negative controls. To date, two works (Flegontov et al. 2015; Sobotka et al. 2017) have investigated the metabolism of chromerids on organellar level, and only the latter work supports the localization of analyzed (plastid) proteins with proteomic data. Our dataset included sequences of typical plastid-targeted proteins as well as proteins with unambiguous localization to mitochondria and other compartments, conserved in other eukaryotic lineages. To ensure that our sequences are largely complete at their N-termini, we used protein models generated by two independent sequencing initiatives, EuPathDB (deposited at CryptoDB; (Woo et al. 2015) and MMETSP (Keeling et al. 2014).

We analyzed the performance of several algorithms based on their sensitivity (percentage of true positive sequences passing a threshold) and precision (percentage of false positives in the set passing the same threshold). For plastid proteomes, ASAFind using a species-specific bit score matrix was found to be the most efficient for each chromerid species. The sensitivity of ASAFind with *Phaeodactylum tricornutum* sequences was comparable to our results (80%; Gruber et al. 2015). With mitochondrial references, we could not observe predictive differences for *V. brassicaformis* sequences. With *C. velia* datasets, HECTAR was more precise at higher sensitivity levels, and MultiLoc2 could not reach an 85% precision (Table 1). TargetP performed similarly to HECTAR (TargetP is indeed part of HECTAR’s mitochondrial module) and is widely used for finding mitochondria-targeted genes with sensitivity around 60-75% (depending on the model and reliability class) in various organisms (Emanuelsson et al. 2007), including the chromerids (Kořený et al. 2011; Flegontov et al. 2015; Woo et al. 2015). This accuracy is also relatively lower because a portion of mitochondrial proteins use alternative routes or signals for translocation to this organelle (Sun & Habermann 2017). Several of our plastid and mitochondrial reference sequences were recovered as false negatives not passing the probability threshold. Indeed, 136 of the 1,141 reference sequences had alternative open reading frames or were unannotated and had to be manually adjusted using homology annotation, alignments and phylogenetic analyses (Supplementary Table S1). For instance, we could not obtain consistent localizations for the enzymes of the MEP pathway that is thought to be exclusively plastid-localized (Fig. 1). With untreated data, the false negative discovery rate would be much higher, leading to orphan enzymes predicted to unexpected compartments. This points out the problems with automated analyses – there is an essential need for highly complete sequence data, which worsens large-scale predictions in the chromerids.

Although targeting presequences are generally not generally not conserved on sequence level, the conservation of amino acids flanking the BTS cleavage sites was found to be crucial for proper plastid targeting (Gruber et al. 2007). Proteins targeted to the rhodophyte-derived complex plastids generally expose an invariant Phe at their N-terminus after signal peptide cleavage (Gould et al. 2006; Patron & Waller 2007). Based on our frequency matrices, in *Vitrella* plastid-targeted proteins the Phe motif is not strictly conserved, while most plastid-targeted proteins of *Chromera* do possess this Phe (Fig. 3). This suggests there is some versatility of the translocation machineries in chromerids. Our observations are similar to the results of Woehle et al. (2011), although the frequency of Phe in their *C. velia* dataset appears higher. AT richness above 57 % was shown to correlate with a shift in amino acid composition of transit peptides towards AT-rich codons (Ralph et al. 2004) but the difference in the GC percentage of the plastid reference transcripts in chromerids appears unlikely to be the cause for a diminished Phe (53.0 % and 59.6 % GC in *C. velia* and *V. brassicaformis*, respectively). Lastly, we cannot exclude the possibility that mis-identified cleavage sites or mis-assembled transcripts in our data affected the amino acid frequencies. Nevertheless, the Phe motif is less strongly retained in *Toxoplasma* and apparently absent in *Plasmodium* apicoplast-targeted proteins (Patron & Waller 2007), consistent with our results. In addition, not all rhodophyte-derived lineages retain a high percentage of plastid-targeted proteins with the Phe; despite Phe occurs predominantly in cryptophytes (*Guillardia theta*) and heterokonts (*Thalassiosira pseudonana* and *Phaeodactylum tricornutum*), haptophytes apparently do not rely on Phe in their transit peptide presequences (Kilian & Kroth 2005; Patron & Waller 2007; Gruber et al. 2015). Phe motif is also absent from the transit peptides of chlorophyte-derived algae (Patron & Waller 2007).

The pathways that localize into plastids are typically associated with photosynthesis. We identified enzymes responsible for the synthesis of tetrapyrroles, terpenoids, carotenoids, lipids, iron-sulfur clusters and carbohydrates, hence compounds required for the proper assembly and function of the photosynthetic machinery. Our findings are consistent with previous biochemical analyses, showing that chromerids have a limited set of photosynthetic pigments (Moore et al. 2008; Kotabová et al. 2011) and that they exhibit unique structural changes to the photosystems (Sobotka et al. 2017). We show that terpenoid and lipid pathways are primed with substrates produced by the plastid carbohydrate metabolism (DHAP, pyruvate, and PEP) and linked with the rate of carbon fixation by substrate availability (Fig. 1). This is not unprecedented, as plant chloroplasts show a similar arrangement in photosynthetically active and inactive plastids (e.g. Neuhaus and Emes 2000).

Nevertheless, plastid products might be essential beyond photosynthesis, which is best illustrated by comparison of algae with relatives that lost photosynthetic abilities (e.g., Hadariová et al. 2018). Indeed, the biology of non-photosynthetic plastids has been best studied in apicomplexans to find suitable weak-spots of these infamous parasites. The apicoplast produces fatty acids and terpenoids and participates in heme synthesis to sustain parasite growth in hosts where salvage of these compounds is not possible (reviewed by van Dooren and Striepen 2013). This dependence on a remnant plastid can be regarded as an evolutionary constraint that cannot be easily overcome once parallel pathways in the cytosol or mitochondria are lost (though losses of plastids occasionally happen, see Füssy & Oborník 2018; Oborník 2018). Consistent with this view, apicomplexans lack apicoplast-independent pathways for terpenoid (via mevalonate), fatty acid (using type I fatty acid synthase, or FASI) and tetrapyrrole synthesis. Some apicomplexans appear to possess FASI-like enzymes, though functional analyses suggest that *Cryptosporidum* FASI is not involved in *de novo* fatty acid synthesis and rather accepts long-chain fatty acyl thioesters as substrates for elongation (Zhu et al. 2004). The importance of *Toxoplasma* FASI remains unclear, while other apicomplexan FASI-like enzymes might be involved in polyketide synthesis (Kohli et al. 2016).

The photosynthetic relatives to apicomplexans, chromerids are also likely to lack cytosolic pathways for fatty acid, terpenoid and tetrapyrrole synthesis and thus rely entirely on the plastid synthesis. In addition, chromerid plastids synthesize FAD cofactors and a nitrogen metabolism intermediate, ornithine (see below). Conversely, chromerid plastids do not host any fatty acid elongases or desaturases, therefore short-chain fatty acids need to be exported, processed and reimported for incorporation into plastid lipids (Botté et al. 2011). This is analogous to the fatty acid synthesis architecture in apicomplexans (Mazumdar & Striepen 2007). Similarly, chromerid tetrapyrrole biosynthesis relies on the import of the starting substrate, ALA, from the cytosol (Kořený et al. 2011). Therefore, chromerid plastids pathways are interdependent with the cytosol, pointing out possible feedback regulatory mechanisms to limit their biosynthetic activity.

In the dark, the reduced triose phosphate supply from the chromerid plastid must be counterbalanced by catabolic reactions of the mitochondrial TCA cycle and respiration. The chromerid TCA cycle and respiratory chain represent modifications to the canonical mitochondrial pathways (see Results). The list of chromerid mitochondrial enzymes is largely shared with parasitic apicomplexans (Flegontov et al. 2015; Jacot et al. 2016). As yet, it is unclear what impact this arrangement has on the metabolism of photoautotrophic organisms, but it stands out that through the TCA cycle chromerid mitochondria are metabolically versatile and integrate catabolic pathways with enzymes priming anabolic reactions and enzymes typically associated with anaerobiosis (Flegontov et al. 2015). Intracellular stages of *Toxoplasma gondii* actively catabolize host glucose via the oxidative TCA cycle to produce energy efficiently (MacRae et al. 2012). In comparison, for asexual stages of *Plasmodium*, purine salvage from oxaloacetate is vital, while the importance of the TCA cycle is diminished (Bulusu et al. 2011; Ke et al. 2015). Similarly, cytosolic ATP citrate lyase contributes to acetyl-CoA production in *T. gondii* (Tymoshenko et al. 2015). As such, chromerid mitochondria are likely to have major influence on metabolic fluxes in the cell.

We observed that amino acid biosynthesis pathways consistently showed cytosolic localizations, with minor exceptions. This is probably not due to mis-annotation as we could not extend the sequences towards an alternative N-terminus in most cases. Chromerid plastids thus appear to host only parts of Cys and Arg synthesis, suggesting that algae do not necessarily synthesize amino acids in plastids as plants do (Van Dingenen et al. 2016). While plastid Cys synthesis is required by the Fe-S cluster assembly system SUF, the plastid production of ornithine for Arg biosynthesis might have a regulatory role in chromerids. Indeed, nitrogen metabolism is energy-demanding and subject to intense cross-talk with carbon metabolism, and plastids play a crucial role in balancing the fluxes (Stitt 2002; Németh et al. 2018). As yet, the primarily cytosolic localization of nitrogen metabolism in chromerids appears extraordinary among phototrophs (Allen et al. 2011; Bromke 2013; de la Torre et al. 2013; Dorrell et al. 2017). Of studied apicomplexans, *Toxoplasma* has the broadest capacity to produce amino acids *de novo* or secondarily from specific precursors, being auxotrophic only for Arg, His and Trp (Chaudhary et al. 2014; Tymoshenko et al. 2015). Although the middle steps of Pro and Lys synthesis are currently unknown (Shanmugasundram et al. 2013), none of these amino acid synthesis pathways appears to be placed exclusively in the apicoplast, in line with our results. Arg scavenging from outer sources might have therefore triggered the loss of the ornithine cycle in the ancestor of apicomplexans. Reliance on amino acid import from the host resulted in even greater reduction of amino acid synthesis capabilities. Thus, *Plasmodium* synthesizes six amino acids (Gly, Glu, Gln, Pro, Asp, Asn), while *Cryptosporidium*, which entirely lacks an apicoplast (Keithly et al. 2000; Chaudhary et al. 2014), synthesizes only Gly, Glu and Pro. It is tempting to speculate that the major role of the cytosol in amino acid synthesis facilitated the plastid loss in a distantly related non-photosynthetic dinoflagellate, *Hematodinium* (Gornik et al. 2015).

In apicomplexans, indeed, the main source of energy is glycolysis of host-drawn glucose, but the parasites appear to use the same carbohydrate subpathways as chromerids. These include the compartment exchange of triose phosphates via phosphate transporters (Lim et al. 2010), parallel cytosolic and plastid glycolysis to pyruvate (Fleige et al. 2007), cytosolic accumulation of storage polysaccharides (Coppin et al. 2005) and a notable relocation of the canonical pyruvate dehydrogenase in the apicoplast to feed the fatty acid synthesis (Foth et al. 2005; Fleige et al. 2007). There are also obvious commonalities in the arrangement of mitochondrial metabolism, including the modifications of the canonical TCA cycle, the respiratory chain, and the anabolic reactions of mitochondria. As a previous comparative analysis of mitochondrial metabolism of apicomplexans and dinoflagellates showed, this arrangement is largely shared among all myzozoans, and likely represents an ancestral state employed by the dinoflagellate-apicomplexan progenitor (Danne et al. 2013).

The overall architecture of biosynthetic pathways in chromerids suggests that their cytosol represents the compartment which integrates the cellular energy status. Through metabolite flow it can directly regulate anabolic and catabolic reactions based on photosynthate supply from the plastid and nutrient availability. Hypothetically, such a central role for the cytosol could have been employed by a free-living unicellular eukaryovore/algivore that ingests its prey into a food vacuole and gradually digests it, much like the one we picture was the ancestor of Myzozoa (Tikhonenkov et al. 2014). Indeed, also dinoflagellates employ a remarkable spectrum of trophic strategies (reviewed in Waller & Kořený 2017). Despite this diversity, there are commonalities in the arrangement of mitochondrial and plastid pathways among myzozoans (Danne et al. 2013; Gornik et al. 2015; Waller et al. 2016), pointing out that the cytosolic pyruvate hub may be persisting through plastid endosymbiotic events.

The apparently ancestral potential to exploit external resources and the richness of apicomplexan cell-surface transporters raises the question whether chromerids also take considerable advantage of extracellular nutrients. Both chromerids were isolated as coral-associated organisms (Moore et al. 2008; Oborník et al. 2012) and intriguing data have been presented on mixotrophy of *C. velia* and its association with corals (Cumbo et al. 2013; Foster et al. 2014; Mohamed et al. 2018). This question therefore needs to be addressed in more detail, as it might present implications on the early evolution of Apicomplexa. However, experimental data are currently unavailable to present an in-depth metabolomic model for chromerids and we can merely compare our results with those from other alveolates. Here we present a benchmark set of biochemical pathways that can be investigated by quantitative or phylogenetic approaches for a deeper understanding of one of the “alveolate ways” of trophic transition from photosynthetic algae to obligatory parasites.

## CONCLUSIONS

Chromerids occupy a pivotal position in the tree of alveolates and hold the key to our understanding of transition to parasitism; they are free-living phototrophic algae with relatively canonical chromosomes and branch sister to the apicomplexans. To sketch a few more pathways on the metabolic map of chromerids, we prepared reference datasets of plastid, cytosolic and mitochondrial enzymes. We have manually curated these protein sequences so that most of them are complete, which is crucial for correct predictions of their subcellular localization. We unveiled that chromerid plastids vividly exchange compounds with the cytosol and mitochondria in order to produce terpenoids, lipids and tetrapyrroles. In contrast, chromerid plastids appear to have only a minor role in amino acid synthesis, as most of these pathways reside in the cytosol. Uniquely, chromerids were found to use a plastid ornithine cycle combined with a cytosolic Arg cycle for synthesis and decomposition of this amino acid. We outlined a major hub represented by lower glycolysis and gluconeogenesis enzymes that appears to regulate carbon and nitrogen metabolite flow depending on the photoactivity of the plastid. When compared to apicomplexan metabolic pathways, our model confirms a conserved architecture of carbon and nitrogen metabolism in these groups. Further analyses are though needed to gain insight into the regulation of these pathways in response to various cues. Using the suggested prediction tools, it is now also possible to introduce more enzymatic steps to the picture.

## ABBREVIATIONS

2-OG: 2-oxoglutarate,
ACP: acyl-carrier protein,
ALA: delta-aminolevulinic acid,
BTS: bipartite (plastid) targeting signal,
Chl *a*: chlorophyll *a*,
DGDG/MGDG: di-/monogalactosyl-diacylglycerolipid,
DHAP: dihydroxyacetone phosphate,
Ery-4P: erythrose 4-phosphate,
FAD: flavin adenine dinucleotide,
GS/GOGAT: glutamine synthase / glutamine oxoglutarate aminotransferase,
IMS: inter-membrane space,
MEP: methyl-D-erythritol 4-phosphate,
mTP: mitochondrial transit peptide,
OAA: oxaloacetate,
orn: ornithine,
PEP: phospho*enol*pyruvate,
PQ: plastoquinol,
PRPP: 5-phosphoribosyl-1-pyrophosphate,
pyr: pyruvate,
Ru-5P: ribulose 5-phosphate,
TCA: tricarboxylic acid,
THF: tetrahydrofolate,
UQ: ubiquinone.

## ETHICS APPROVAL AND CONSENT TO PARTICIPATE

Not applicable.

## CONSENT FOR PUBLICATION

Not applicable.

## AUTHORS’ CONTRIBUTIONS

MO and ZF conceived the study. ZF and TF performed all the bioinformatic analyses and prepared the figures and tables. ZF drafted the manuscript. MO acquired funding. All authors edited the manuscript and approved its final form.

### ACKNOWLEDGEMENTS

The authors would like to thank the Czech Science Foundation (project 16-24027S granted to MO) and the European Regional Development Fund (ERDF) (No. CZ.02.1.01/0.0/0.0/16_019/0000759) for funding. Computational resources from MetaCentrum and CERIT-SC, Brno, Czech Republic, are greatly appreciated. We are grateful to Ansgar Gruber for helpful discussions.

## SUPPLEMENTARY MATERIAL

**Supplementary Table S1: Reference protein lists grouped by metabolic function; inferred phylogenies of argJ, polyprenyl-PP synthases, and PRPP synthase; set of updated reference protein sequences.** Trees in Newick format, accessions are marked on leaves. The maximum-likelihood trees were inferred by the IQ-TREE software (see main text Methods). Sequence updates are based on homology searches within alternative transcriptomic datasets (elongated sequences) and comparison of orthologous sequences in *C. velia* and *V. brassicaformis* (merged contigs).

**Supplementary Table S2: Prediction scores for reference proteins and the species-specific ASAFind scoring matrices of C. velia and V. brassicaformis.**

**Supplementary Figure S3: Targeting peptide score distributions for reference sequences.** Reference colors: green – plastid, blue – mitochondrial, grey – other (cytosolic or non-plastid endomembrane route).

